# A single-cell atlas of *de novo* β-cell regeneration reveals the contribution of hybrid β/δ-cells to diabetes recovery in zebrafish

**DOI:** 10.1101/2021.06.24.449704

**Authors:** Sumeet Pal Singh, Prateek Chawla, Alisa Hnatiuk, Margrit Kamel, Luis Delgadillo Silva, Bastiaan Spanjard, Sema Elif Eski, Sharan Janjuha, Pedro Olivares, Oezge Kayisoglu, Fabian Rost, Juliane Bläsche, Annekathrin Kränkel, Andreas Petzold, Thomas Kurth, Susanne Reinhardt, Jan Philipp Junker, Nikolay Ninov

## Abstract

Regeneration-competent species possess the ability to reverse the progression of severe diseases by restoring the function of the damaged tissue. However, the cellular dynamics underlying this capability remain unexplored. Here, we use single-cell transcriptomics to map *de novo* β-cell regeneration during induction and recovery from diabetes in zebrafish. We show that the zebrafish has evolved two distinct types of somatostatin-producing δ-cells, which we term δ1- and δ2-cells. Moreover, we characterize a small population of glucose-responsive islet cells, which share the hormones and fate-determinants of both β- and δ1-cells. The transcriptomic analysis of β-cell regeneration reveals that β/δ hybrid cells constitute a prominent source of insulin-expression during diabetes recovery. Using *in vivo* calcium imaging and cell tracking, we further show that the hybrid cells form *de novo* and acquire glucose-responsiveness in the course of regeneration. The overexpression of *dkk3*, a gene enriched in hybrid cells, increases their formation in the absence of β-cell injury. Finally, interspecies comparison shows that plastic δ1-cells are partially related to PP-cells in the human pancreas. Our work provides an atlas of β-cell regeneration and indicates that the rapid formation of glucose-responsive hybrid cells contributes to the resolution of diabetes in zebrafish

## Introduction

The pancreas, a glandular organ, is composed of exocrine and endocrine compartments. The exocrine part of the pancreas is involved in food digestion via the function of acinar and ductal cells. The endocrine part is responsible for metabolic homeostasis, specifically regulation of blood glucose levels. The endocrine compartment is organized into the islets of Langerhans, which consist of three major cell types: α-, β- and δ-cells. The three cell-types produce and secrete specific hormones: α-cells generate glucagon (*gcg*), β-cells generate insulin (*ins*), and δ-cells generate somatostatin (*sst*). Blood glucose homeostasis is achieved by regulated secretion of endocrine hormones in the blood stream; insulin activates glucose uptake in peripheral tissue, while glucagon opposes insulin action by stimulating the production of sugar in the liver (Noguchi and Huising, 2019). Somatostatin release has an inhibitory role for both insulin and glucagon secretion, as its loss triggers acute hypoglycemia (Li et al., 2018). As the selective autoimmune destruction of β-cells causes Type 1 Diabetes, enhancing the limited regenerative potential of the pancreas has become an important quest towards developing new regenerative therapies.

Regeneration and cell plasticity are intimately connected. Upon near-complete β-cell destruction, a small proportion of glucagon-expressing α-cells trans-differentiate into β-cells in adult mice and embryonic zebrafish (Chera et al., 2014; Thorel et al., 2010). In addition, a subset of terminal duct cells, called centroacinar cells can give raise to β-cells in adult fish and mice (Delaspre et al., 2015; Mameishvili et al., 2019). Moreover, in juvenile mice, δ-cells initiate insulin production upon ablation of the β-cells, however this cellular makeover ceases to operate in older animals. To define the full spectrum of cellular dynamics underlying β-cell regeneration requires technologies that can track gene expression in multiple cells over time during the process of β-cells regeneration.

In this study, we applied a unique blend of *in vivo* imaging, single-cell genomics, and genetics to study *de novo* β-cell regeneration in zebrafish. While regeneration is incomplete in mammals, zebrafish can naturally recover from extreme β-cell destruction and hyperglycemia (Moss et al., 2009). However, the reasons underlying this ability remain unexplored. By categorizing the cells in the zebrafish pancreas, we identify two discrete populations of δ-cells, which express different *sst* paralogues. Moreover, we find that a subset of islet cells exist under a hybrid identity, sharing the hormones and fate-determinants of both δ- and β-cells. During ablation and subsequent regeneration, the hybrid cells form *de novo*, acquire glucose-responsiveness and serve as a major source of insulin-expression. We also show that the secreted protein Dkk3b can increase the formation of hybrid cells in the absence of injury. We propose that the rapid formation of glucose-responsive hybrid cells provides a shortcut to restoring glucose homeostasis and contributes to the ability of zebrafish to reverse the course of diabetes.

## Results

### Single-cell RNA-sequencing of the pancreas identifies two populations of δ-cells in zebrafish

We first characterized the cell types in the zebrafish pancreas under homeostasis. We analyzed the data from a droplet based single-cell transcriptome profiling of the islets and surrounding exocrine tissues from 2 months post-fertilization (mpf) zebrafish (Salem et al., 2019). For this experiment, the tissue surrounding the principal islet was dissected from the pancreas (n = 6 animals). The dissected tissue was enzymatically dissociated, sorted by flow cytometry to remove damaged cells and profiled using the 10x Genomics pipeline (Fig. 1A). The transcriptome data was subjected to quality control that included number of unique molecular identifiers (UMI), genes detected per cell, and the percentage of mitochondrial reads detected per cell. This yielded a total of 1969 cells for downstream analysis. Clustering of the single-cell transcriptome data using Seurat identified eight major clusters in the data (Fig. 1B-D). Based on the expression of marker genes, the clusters corresponded to acinar (*ela2*), ductal (*cftr, hnf1ba, her15.1, anxa4;* Fig. S1A), endothelial (*vsg1*, also known as *plvapb*), β-cells (*ins*), α-cells (*gcgb*), ͼ-cells (*ghrl*), and two discrete types of δ-cells (*sst1.1* and *sst2*) (Fig. 1B-C) (Table S1). The two groups of δ-cells include a population marked by *sst2 (δ2)* (Li et al., 2009) and a distinct type that expresses *sst1.1 (δ1)*. Each of the two *sst* genes, which arose through gene-duplication in teleost fish is predicted to encode for a full-length and functional somatostatin protein (Devos et al., 2002; Liu et al., 2010). One notable difference between the δ1- and δ2-cell types is the preferential expression *of pancreatic and duodenal homeobox 1* (*pdx1)* in the former group (Fig. 2A). A number of other genes were differentially expressed between the two δ-cell populations (Fig. 2B and Table S2). Thus, the zebrafish islet is equipped with two distinct δ-cell types, each expressing a different *sst* paralogue.

**Figure 1.**
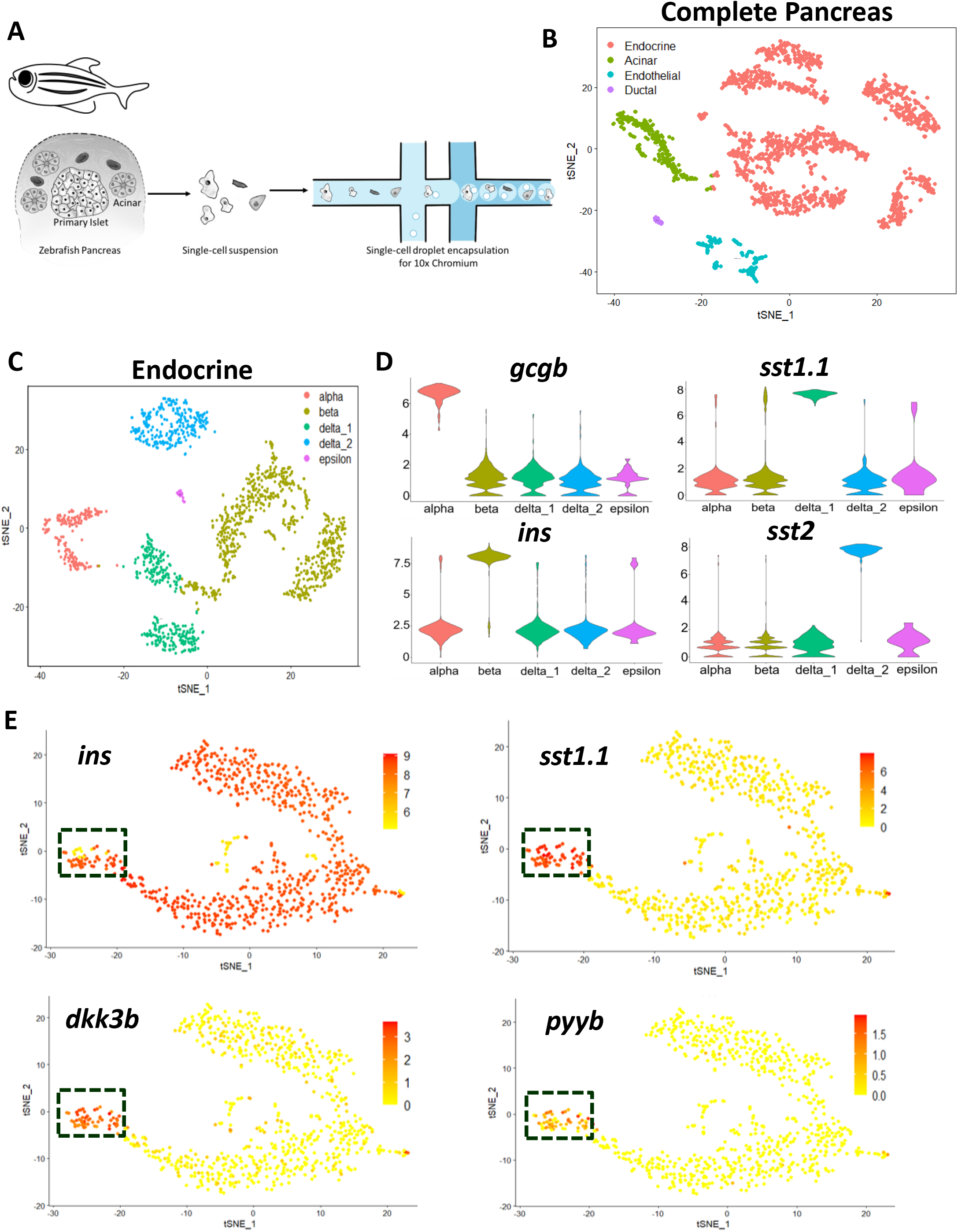
Single-cell RNA-sequencing of the zebrafish pancreas reveals two types of islet δ-cells. **(A)** Schematic of the high-throughput 10x Droplet-Seq. approach to classify the cell populations in the pancreas from 2 mpf zebrafish. **(B)** t-SNE plot representation of 1,969 pancreatic cells with clusters representing distinct cell-types. **(C)** t-SNE plot representation of 1,530 endocrine cells with clusters based on the expression of hormonal genes. **(D)** Violin Plot for expression of hormonal genes in the endocrine clusters. **(E)** t-SNE plot representation of 746 insulin-expressing pancreatic cells with specific gene expression overlaid as a heat-map. Cells with co-expression of *insulin* and *sst1.1* are outlined.

**Figure 2.**
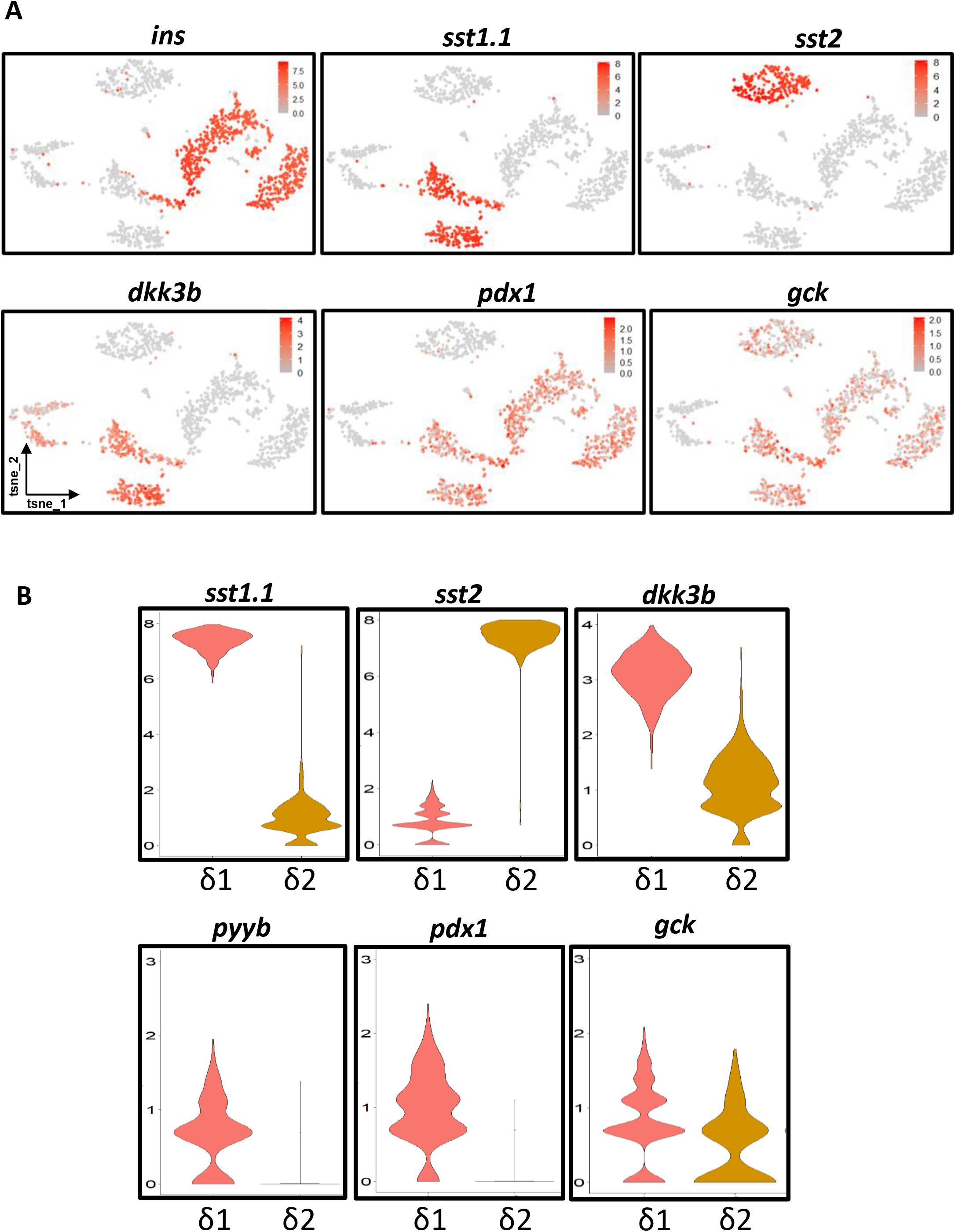
Markers of δ1- and δ2-cells. **(A)** t-SNE plot representation of the β-, δ1- and δ2-cell populations, expressing *ins*, *sst1.1* and *sst2*, respectively. glucokinase (*gck*) is expressed in both δ1- and δ2-cells, whereas *dkk3b* and *pdx1* are enriched in the δ1-cells. **(B)** Violin plot highlighting the expression levels of *sst1.1*, *sst2*, *dkk3b*, *pyyb, pdx1* and *gck* in the δ1- and δ2-cell clusters.

### The zebrafish islets contain a persistent bi-hormonal cell population

Most cell populations we identified contained discrete hormonal cell-types. One exception occurs in the cells of the *ins-* and *sst1.1*-lineages. Notably, ∼8 % of all *ins*-expressing cells (60 cells out of 746) were positive for both *ins* and *sst1.1* (Fig. 1E). The bi-hormonal cells had further distinguishing features. The cells expressed *dkk3b*, a Dickkopf-related secreted protein, and *pyyb*, a short peptide related to Protein YY (Fig. 1E). *dkk3b* and *pyyb* were also enriched in δ1-cells compared to δ2-cells (Fig. 2).

To confirm the co-expression of *sst1.1* and *ins*, we utilized the single-cell transcriptomic profiles of sorted β-cells from seven stages of zebrafish life including the juvenile and adult periods (Singh et al., 2018). We detected above-background expression of *sst1.1* in a fraction of *ins*-expressing cells at each of the seven stages (Fig. S2A). The fraction of *sst1.1*-expressing cells ranged from 5 to 9 % of the *ins*-positive population throughout the life of the fish, showing that the bi-hormonal population maintains steady proportions.

Using Assay for Transposase-Accessible Chromatin (ATAC) of sorted β-cells from 2-mpf zebrafish, we detected open chromatin peaks in regulatory elements of *ins*, *sst1.1* and *dkk3b* (Fig. S2B). This suggests that *ins*-expressing cells have transcriptionally favorable chromatin for expression of genes enriched in the bi-hormonal cells.

### Histological assessment of the bi-hormonal cells in the zebrafish pancreas

To directly visualize the bi-hormonal cells in the zebrafish pancreas, we took advantage of a transgenic reporter line, *Tg(sst1.1: EGFP-Ras)* (Löhr et al., 2018) in which the regulatory sequences of the *sst1.1* gene drive a membrane localized green fluorescent protein (Fig. 3A). Within islets, the GFP-positive δ1-cells formed an intricate mesh of cells distributed throughout the zebrafish islet and are interconnected by long cellular processes (Figure 3B and Fig. S3A). We stained the pancreas from *Tg(sst1.1: EGFP-Ras)* larvae with antibodies against insulin to mark the β-cells. Confocal imaging of the islets demonstrated the presence of individual cells that co-expressed both EGFP and insulin (Fig. 3B). We further confirmed the co-expression of somatostatin and insulin proteins using immunofluorescence in pancreatic sections from adult fish aged 7 and 16 mpf (Fig. 3C-D). We clearly detected immuno-reactivity for the two hormones along with the membrane GFP maker in single cells, corroborating the presence of bi-hormonal cells in the islet. In the bi-hormonal cells, insulin showed higher concentration in the cytoplasm whereas somatostatin was localized closer to the cell membrane, which may reflect differential distribution of secretary granules for the two hormones. To investigate if the ins/sst1.1 bi-hormonal cells occupy a specific position in the islet, we determined their distribution in adult zebrafish islets. We observed that the bi-hormonal cells are predominantly present in the islet’s periphery with fewer cells in the center (Fig S3B). Based on these results, along with the single-cell transcriptomic data, we have identified cells that express at the RNA and protein levels both insulin and somatostatin and that persist from the early stages of development to adulthood.

**Figure 3.**
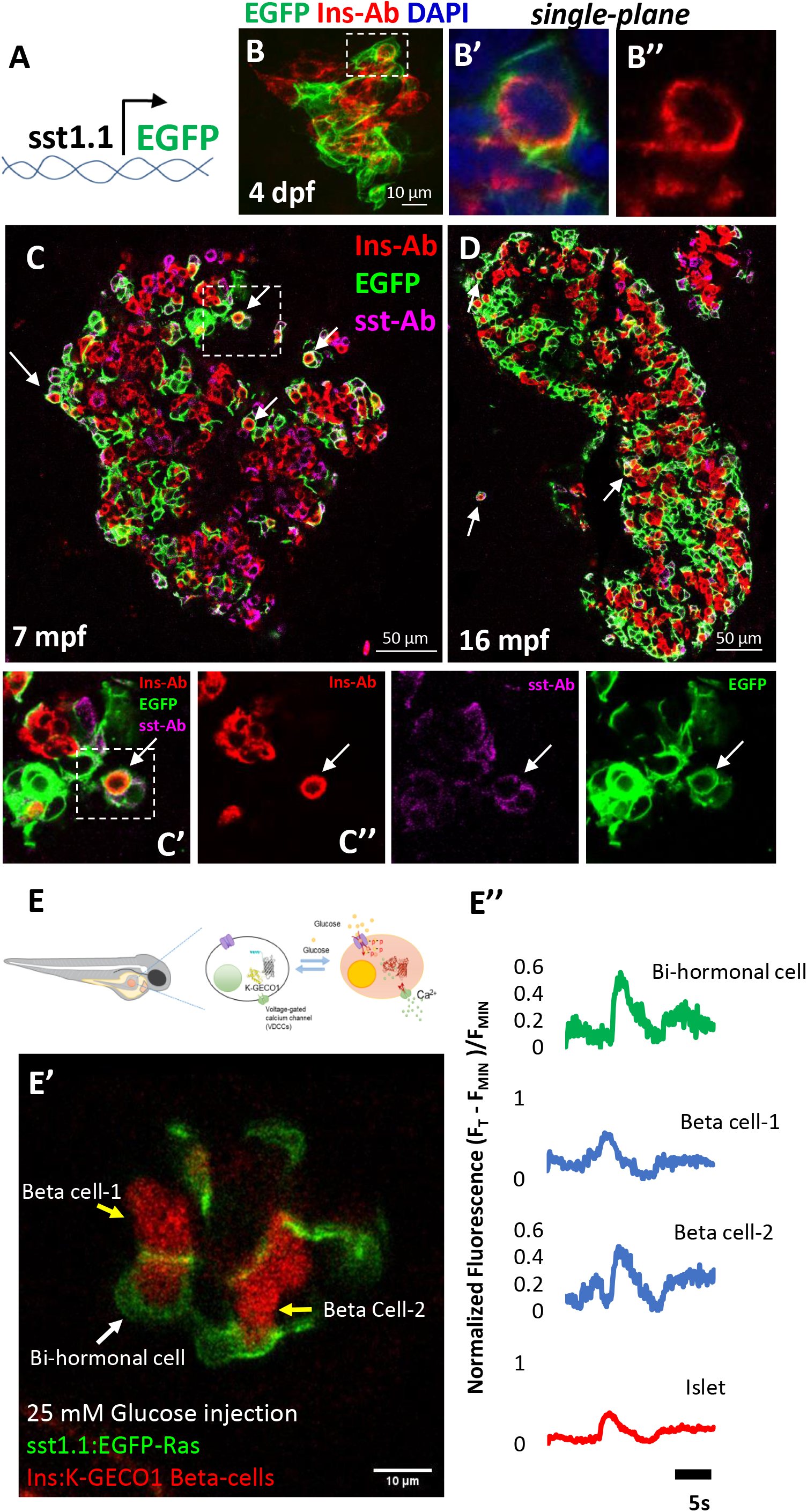
Characterization of a small population of glucose-responsive β/δ1 hybrid cells. **(A)** Schematic of the *Tg(sst1.1:EGFP-Ras)* reporter line used to mark specifically the *sst1.1* population, where a BAC containing the *sst1.1* promoter drives membrane GFP expression. **(B)** Confocal projection of whole-mounted islets from the *Tg(sst1.1: EGFP-Ras)* reporter line stained with an anti-insulin antibody. The dashed line indicates a GFP and insulin co-expressing cell. The high-magnification images in B’ and B’’ shows a single confocal plane of the same cell co-expressing GFP and insulin. **(C-D)** Thin pancreatic sections from the *Tg(sst1.1: EGFP-Ras)* reporter line at 7 months post-fertilization (mpf) and 16 mpf. The sections were stained with insulin (red) and somatostatin (magenta) antibodies. Arrows mark EGFP+ cells co-stained with insulin and somatostatin antibody. The white box indicates the region displayed at higher magnification in C, revealing cells expressing each protein along with GFP. **(E-E’’)** Two-Photon in-vivo Ca^2+^ imaging in 5 dpf double-transgenic zebrafish larvae of the genotype *Tg(sst1.1: EGFP-Ras); Tg(ins: K-GECO1)* showing the glucose-stimulated response of mono-hormonal and bi-hormonal cells. Normalized *K-GECO* fluorescent traces, showing glucose-stimulated calcium influx of bi-hormonal and mono-hormonal β-cells.

### The bi-hormonal cells exhibit glucose-stimulated calcium influx *in vivo*

In order to investigate if the bi-hormonal cells are glucose-responsive and if they present influx of calcium ions upon glucose stimulation, we performed 2-Photon calcium imaging *in vivo* using *Tg(sst1.1:EGFP-Ras); Tg(ins:K-GECO1)* double transgenic larvae. We used our previously developed method for calcium imaging (Salem et al., 2019). We found that cells co-expressing EGFP and K-GECO1 show glucose-stimulated calcium influx, which was comparable to the response of β-cells in terms of speed and amplitude of response (n=3 islets) (Fig. 3E). Thus, the hybrid cells are functionally similar to β-cells in their capacity to respond to glucose.

### The bi-hormonal cells are molecular hybrids between δ1- and β-cells

To characterize the bi-hormonal cells at a better molecular resolution than the one provided by the 10x Genomics pipeline, we collected pancreatic tissue from 2 mpf old fish and separated using index flow cytometry β-, δ1-cell and hybrid cells based on the expression of *ins:*tdTomato and *sst1.1:*EGFP-Ras (Fig. 4A). We performed single-cell transcriptomics using the Smart-Seq2 protocol, which has higher sensitivity than droplet-based single-cell sequencing (Ziegenhain et al., 2017). The single-cell sequencing revealed enrichment of *sst1.1* and *ins* in δ1-cells and β-cells, respectively, validating the utility of the reporter lines to distinguish the two cell populations (Fig. 4B-C). Further, we observed a strong Pearson correlation in the co-expression of *sst1.1* and *dkk3b* as well as of *sst1.1* and *pyyb* in individual cells (Fig. 4D), validating our previous observations. Differential gene expression analysis showed that β*-*cells express significantly higher (p_adj_ < 0.05) levels of *nkx6.2* and *mnx1* (Fig. 4E), genes involved in their specification and maturation (Binot et al., 2010; Pan et al., 2015). Conversely, δ1-cells expressed *hhex* (Fig. 4E), a transcription factor involved in the maintenance of δ-cell identity (Zhang et al., 2014). Interestingly, we observed intermediate expression levels for *nkx6.2*, *mnx1* and *hhex* in the bi-hormonal cells, revealing an overlap of β- and δ-cell identity genes within the mixed population.

**Figure 4.**
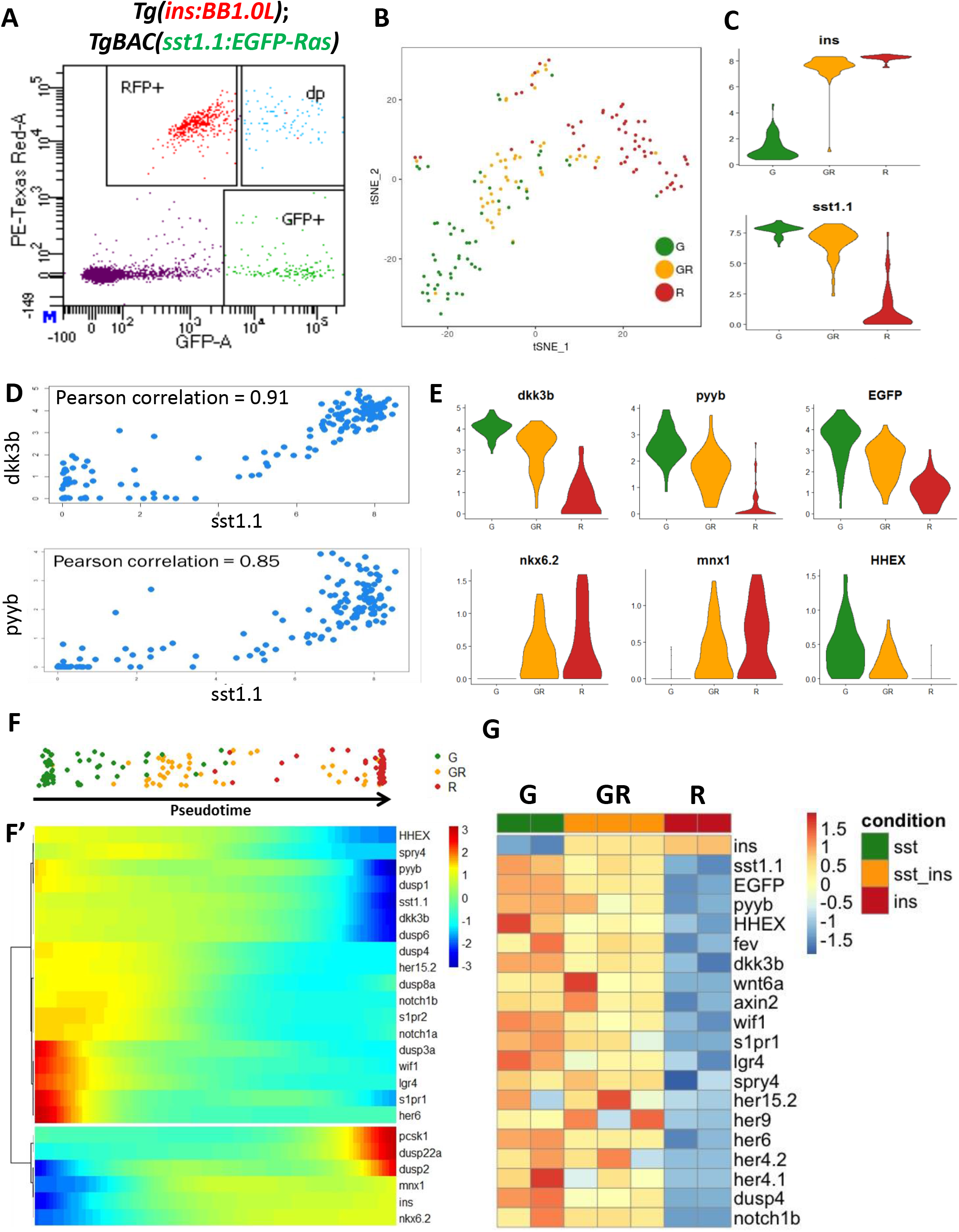
Molecular analysis of the hybrid cells reveals a progenitor signature. **(A)** FACS plot of cells from *Tg(ins:BB1.0L)*; *Tg(sst1.1: EGFP-Ras)* adult fish. Cells were marked based on the presence of red and green fluorescence, and index-sorted into 96-well plates. β-cells are RFP+ (R), δ1-cells are GFP+ (G) and bi-hormonal cells are green+ and red+ (GR). **(B)** A t-sne plot of the profiled cells with colors corresponding to their respective location on the FACS plot. The cells are segregated into three groups: R (β-cells), G (δ1-cells), and GR (bi-hormonal cells). **(C)** Violin Plot showing *insulin* and *sst1.1* expression in the three cell populations. The Y-axis represents normalized gene expression. **(D)** A dot-plot displaying the Pearson correlation coefficient between *sst1.1* and *dkk3b*, as well as, between *sst1.1* and *pyyb* expression levels. **(E)** Violin Plot showing the expression levels of control genes (*dkk3b*, *pyyb*, *EGFP*) and selected genes related to β-cell differentiation and function in the three populations. *mnx1* and *nkx6.2* are enriched in β-cells, whereas *hhex* is expressed in δ1-cells. The Y-axis represents normalized gene expression. **(F)** Pseudo-temporal ordering of the cells, with (F’) showing the expression trend of selected genes that show significant correlation with the pseudo-time. The δ1-cells show enrichment of genes involved in the Wnt, FGF, Notch, and sphingolipid signaling pathways compared to β-cells. The bi-hormonal cells exhibit intermediate expression of genes involved in β- and δ-cell fate-specification and function compared to the pure cell populations. **(G)** To validate the single-cell RNA-Seq. results, the three endocrine cell populations were profiled using bulk-RNA sequencing. The results from the experiment are depicted as a heat-map demonstrating the expression levels of selected genes including those involved in Wnt, FGF and Notch and sphingolipid signaling. All depicted genes show a statistically significant difference in expression between the δ1 and β-cells.

### The bi-hormonal cells express endocrine-progenitor markers

Next, we applied pseudo-temporal analysis using Monocle (Trapnell et al., 2014) to model cellular transitions. The pseudo-temporal model showed that the bi-hormonal cells were positioned in between β- and δ1-cells (Fig. 4F), confirming the hybrid signature of these cells. We further characterized gene expression dynamics by calculating expression changes along the pseudotime (Fig. 4F’). As expected, transcription factors implicated in β-cell lineage and identity (*nkx6.2*, *mnx1*) were enriched towards the β-side of the pseudotime. Moreover, *proprotein convertase subtilisin/kexin type 1* (*pcsk1*) (Benjannet et al., 1991), a gene necessary for conversion of pre-proinsulin to functional insulin hormone displayed a similar trend. In contrast, genes implicated in δ-cell identity (*hhex*), along with pathways controlling endocrine progenitor maintenance such a Wnt, Fgf, Notch, and Sphingolipid signaling (*s1pr1* and *s1pr2*) were associated with the trajectory towards the δ1-cells (Ninov et al., 2012; Serafimidis et al., 2017; Seymour et al., 2012; Sharon et al., 2019; Zhang et al., 2014). The hybrid cells expressed intermediate levels of pathway components related to these pathways, such as *dusp3a, wif1, lgr4, s1pr1, and her6*, as compared to either β- or δ1-cells. The expression of the pathway components involved in progenitor maintenance was confirmed by bulk RNA-sequencing of the sorted cells (Fig. 4G), which also revealed enrichment of *fev*, a transcription factor that was recently shown to be expressed in endocrine progenitors in mice (Byrnes et al., 2018). Altogether, we conclude that the hybrid cells display the transcriptional profiles of intermediate cell states between δ1- and β-cells, and exhibit an enrichment of pathways associated with endocrine progenitors.

### The hybrid cells increase during β-cell regeneration

Intrigued by the association of precursor characteristics of hybrid cells, we further characterized their behaviors during β-cell regeneration in larval zebrafish. Near-complete β-cell death was induced using the nitroreductase (NTR) system (Curado et al., 2007; Pisharath et al., 2007). In this system, *Tg(ins:FLAG-NTR)* transgenic zebrafish, expressing NTR (nitroreductase) specifically in the β-cells, are exposed to Mtz (metronidazole). NTR converts Mtz to a cytotoxin, which specifically kills the β-cells while sparing all other cells in the body. To label *sst1.1*-expressing cells during β-cell regeneration, we utilized *Tg(ins:FLAG-NTR)*; *Tg(sst1.1:EGFP-Ras)* double transgenic line. We exposed *Tg(ins:FLAG-NTR)* larvae to Mtz for 24 hours, and collected samples for observation 4 and 8 days post ablation (dpa) (Fig. S4A). At both stages, we observed a dramatic increase in the percentage of hybrid cells within the regenerating islet (Fig. S4B,C). Of note, when we examined the expression of *sst2*:RFP, which marks the population of δ2-cells (δ2) in zebrafish (Li et al., 2009), we did not observe the formation of hybrid cells (O.K., unpublished),

### A single-cell map of adult β-cell regeneration reveals a major contribution of hybrid cells to insulin expression

To assess the dynamics of hybrid cells during the more physiologically-relevant setting of adult β-cell regeneration, we performed single-cell RNA sequencing (using the 10X genomics pipeline) at different times following β-cell destruction and regeneration in adult zebrafish (9 mpf of age). We selectively ablated the β-cells and dissected the primary islets and surrounding tissues, including the exocrine cells from adult zebrafish at 0, 2, 7, and 14 dpa. These time-points were chosen as they include the time of β-cell destruction, subsequent hyperglycemia, followed by normalization of glucose levels after two weeks of regeneration (Fig. S5) (Delaspre et al., 2015). To induce efficient ablation of insulin-expressing cells, we used a new ablation line, which employs an improved and more effective version of the NTR enzyme (Fig. S5) (Singh et al., 2017). In the unablated samples, we identified all major cell types, including discrete populations of *ins*-expressing β-cells, *sst1.1*-expressing δ-cells, *sst2*-expressing δ2-cells, ductal and acinar cells (Fig. 5 and Table S3). Shortly upon β-cell ablation (at 0 and 2 dpa), the *ins*-only expressing population observed in controls became nearly-completely absent based on single-cell gene expression analysis. Instead, *ins* expression was detected mainly within the δ1-population, in which cells co-expressed both *ins* and *sst1.1* (Fig. 5 and Fig. S6). At 7 and 14 dpa, the hybrid cells increased further due to greater proportions of *in*s-expressing cells in the cluster corresponding to δ1-cells (Fig. S7). Clustering of the integrated data from all time-points revealed that the hybrid cells within the *sst1.1* cluster are the main source of insulin expression during regeneration (Fig. 5D,E). Moreover, upon β-cell destruction, we detected increased expression in *pcsk1* and *pcsk2* in the *sst1.1*-expressing cells, which are necessary for insulin maturation (Fig. S8). These changes in gene-expression were accompanied by an increase in the number of *sst1.1-*expressing cells relative to ablation-invariant α-cells (Fig. S9). At 7 dpa, the relative number of *sst1.1*-expressing cells has stabilized at roughly twice as many as in controls.

**Figure 5.**
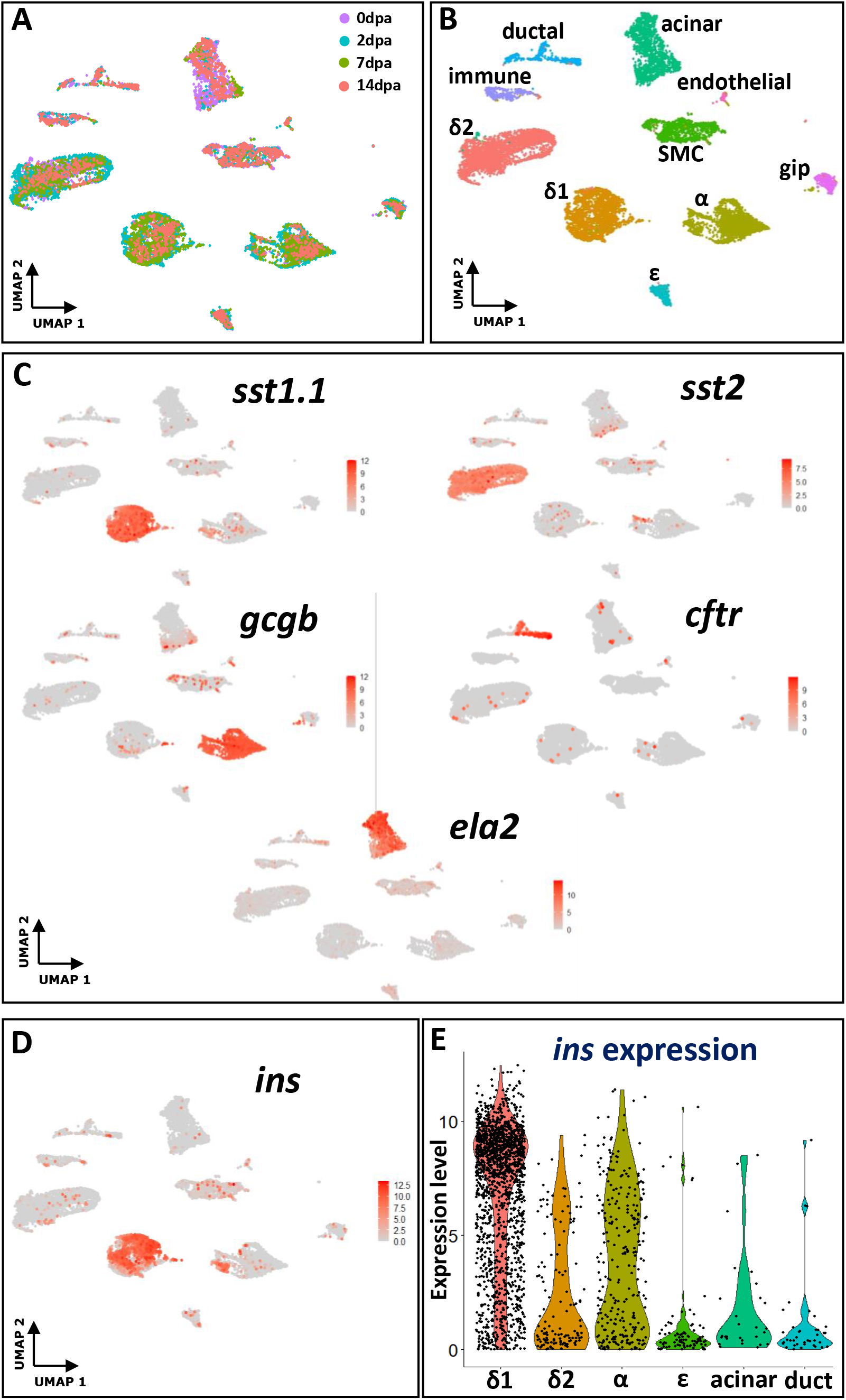
The hybrid cells constitute a prominent source of insulin expression during diabetes recovery in adult zebrafish. **(A-B)** UMAP-plot representation of integrated 10X scRNASeq data from all β-cell regeneration time-points: 0, 2, 7 and 14 days post-ablation (dpa) showing distinct clusters for δ1, δ2, alpha, acinar and duct cells. **(C)** Feature-plot highlighting the expression of *sst1.1 (δ1)*, *sst2 (δ2)*, *gcgb (alpha)*, *cftr (duct)* and *ela2 (acinar)* genes. **(D)** Feature-plot highlighting *ins* expression in different pancreatic cells during regeneration **(E)** Vlnplot highlighting *ins* expression in different pancreatic cells. The β/δ1 hybrid cells among the δ1-cells serve as the main source of insulin-expression during the course of regeneration.

Compared to the *sst1.1*-expressing cells, δ2- and α-cells, showed a relatively minor contribution to insulin expression (Fig. 5D, E). A closer look at the α-cell cluster highlighted that some *gcgb*-expressing cells do show low-to-medium *ins* expression over the course of β-cell regeneration (Fig. S10A-D). This suggests that individual α-cells also act as sources of insulin expression. However, compared to the δ1-cells, α-cells are not the major contributors. Finally, we generated a list of differentially expressed genes (DEGs) in α/β and δ/β hybrid cells at 7 dpa and performed Gene Ontology (GO) analysis (Table S4). Whereas δ/β hybrid cells showed an enrichment of genes involved in biological processes including “type B pancreatic cell differentiation”, “activation of MAPKKK activity” and “negative regulation of canonical Wnt signaling”, the α/β hybrid cells were associated with genes linked to “fatty acid elongation”, “protein peptidyl-prolyl isomerization”, “positive regulation of cytokine production and angiogenesis” and “neuropeptide signaling pathway”. This analysis hints that different molecular mechanisms underlie the formation of α/β and δ/β hybrid cells (Fig. S10E).

### The hybrid cells retain bi-hormonal characteristics

To investigate if δ/β hybrid cells may resolve into *sst1.1+* and *ins+* populations during regeneration, we first analyzed the single-cell transcriptome data from 7 and 14 dpa (Fig. 6). 7 dpa represents an intermediate stage while 14 dpa corresponds to the resolution of diabetes. By specifically focusing on cells that express the two hormones, we observed overlap between the *sst1.1*+ and *ins*+ expressing cells at 7 dpa (Fig. 6A,B). By contrast, at 14 dpa the cells segregated into two clusters (Fig. 6D,E). Notably, the two clusters observed at 14 dpa were enriched for the expression of either *sst1.1* or *ins* (Fig. 6E), indicating that the two cell populations are diverging. We further performed gene correlation analysis to examine the distribution of *sst1.1* and *ins* gene expression level at cellular resolution. At 7 dpa, a positive correlation was observed between *ins* and *sst1.1*-expression (Fig. 6C), suggesting the presence of hybrid cells at this stage. At 14 dpa, however, individual cells displayed a negative correlation between *sst1.1* and *ins-*expression (Fig. 6F), suggesting a trend towards the resolution of gene expression into β- or δ1-cells.

**Figure 6.**
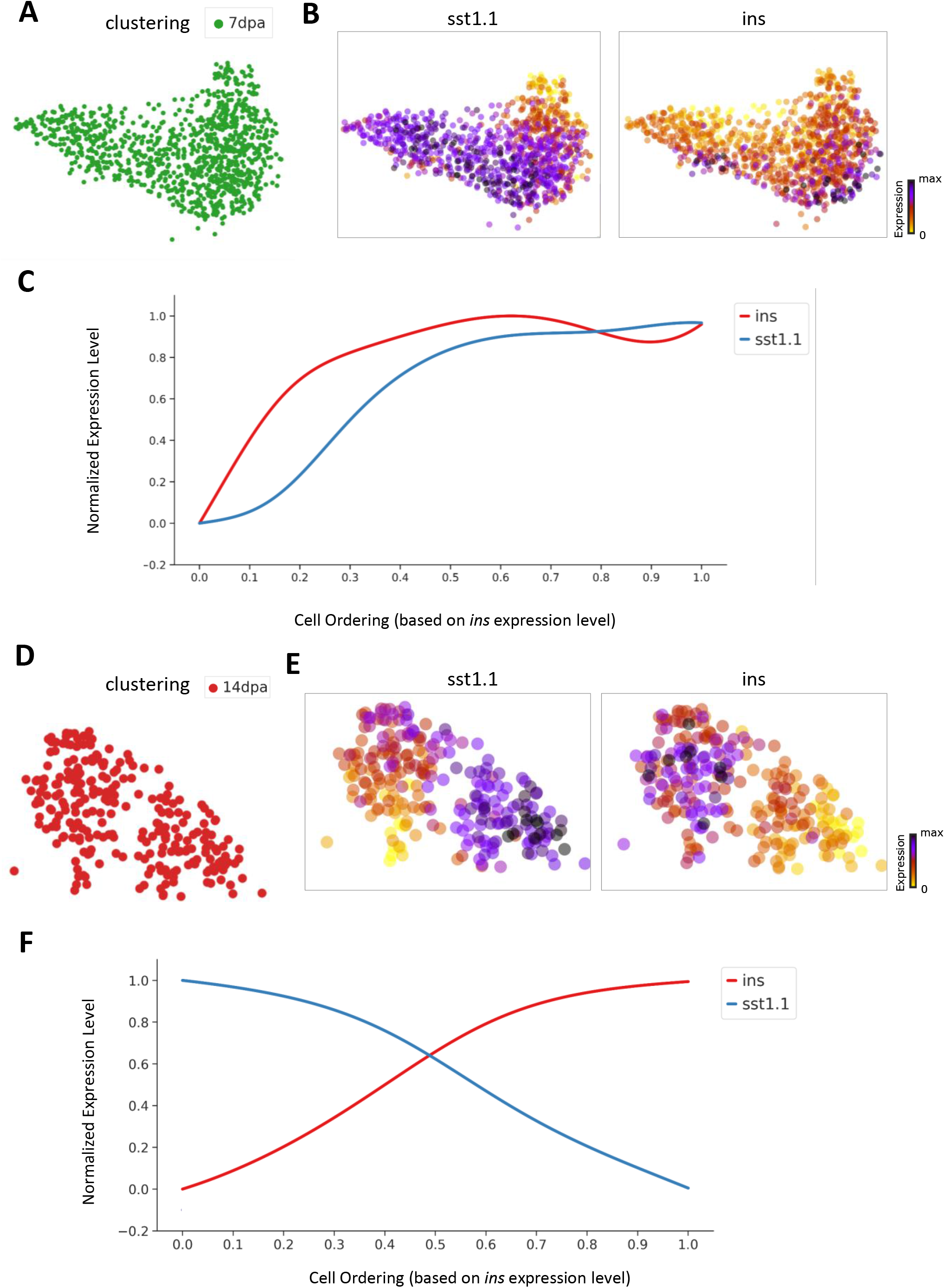
Gene-correlation analysis indicates resolution of the hybrid cells towards *ins*+ and *sst1.1*+ populations during regeneration. (**A**, **D**) UMAP displaying selected cells from 7 dpa (A) and 14 dpa (F). (**B**-**E**) UMAP overlaid with gene expression plots for *sst1.1* and *ins* at 7 dpa (B) and 14 dpa (E). (**C**,**F**) Expression trends for *sst1.1* and *ins* at 7 dpa (C) and 14 dpa (F). At 14 dpa, individual cells along display a negative correlation between *sst1.1* and *ins* expression, whereas at 7 dpa the cells co-express both hormones.

To corroborate our observations from the transcriptome analysis, we co-stained the adult zebrafish pancreatic tissues sections with somatostatin and insulin antibody. Compared to the control samples where cells distinctly expressed insulin and somatostatin, we observed a near-to-complete loss of insulin-positive-cells at 2 dpa (Fig. S11A, B). At 7 and 14 dpa, we found that the insulin expressing cells co-expressed somatostatin. Notably, the majority of the insulin-positive cells continued to exhibit bi-hormonal signature until 14 dpa (Fig. S11C, D). Thereafter, we also characterized the secondary islets over the course of regeneration. Similar to the primary islets, the secondary islets in the unablated samples did not show a significant overlap between the somatostatin and insulin-positive cells. However, there was a predominance of ins/sst bi-hormonal cells both at 7 dpa and at 14 dpa (Fig. S12). Thus, the immunohistological results contrast with the transcriptomics analysis and indicate that the resolution of the cells into mono-hormonal cells is incomplete until at least 2 weeks post-β-cell loss.

### Live-cell tracking reveals hybrid-cell transition during regeneration

To assess if the bi-hormonal cells arise *de novo* during β-cell regeneration or whether they might reflect pre-existing cells that have survived the ablation procedure, we performed live imaging of islets undergoing regeneration during the optically translucent larval stage. Insulin-expressing cells were labeled using *ins:*tdTomato, whereas *sst1.1*-expressing cells were marked using *sst1.1:*EGFP-Ras. Ablation of β-cells was induced using *Tg(ins:FLAG-NTR)* by Mtz-treatment from 3 to 4 dpf. First, we measured the efficiency of cell ablation and found that both β-cells and hybrid cells were efficiently ablated (Fig. 7A). We then monitored longitudinally the regeneration process in individual larvae. Strikingly, δ1-cells showed dramatic changes in morphology, as they accumulated prominent membrane-bound vacuolar structures within their cytosol evident both under confocal and electron microscopy (Fig. S13). Moreover, within islets that previously had lost any tdTomato expression, we now observed its re-expression in EGFP-positive cells, indicating the *de novo* origin of the hybrid cells (Fig. 7A).

**Figure 7.**
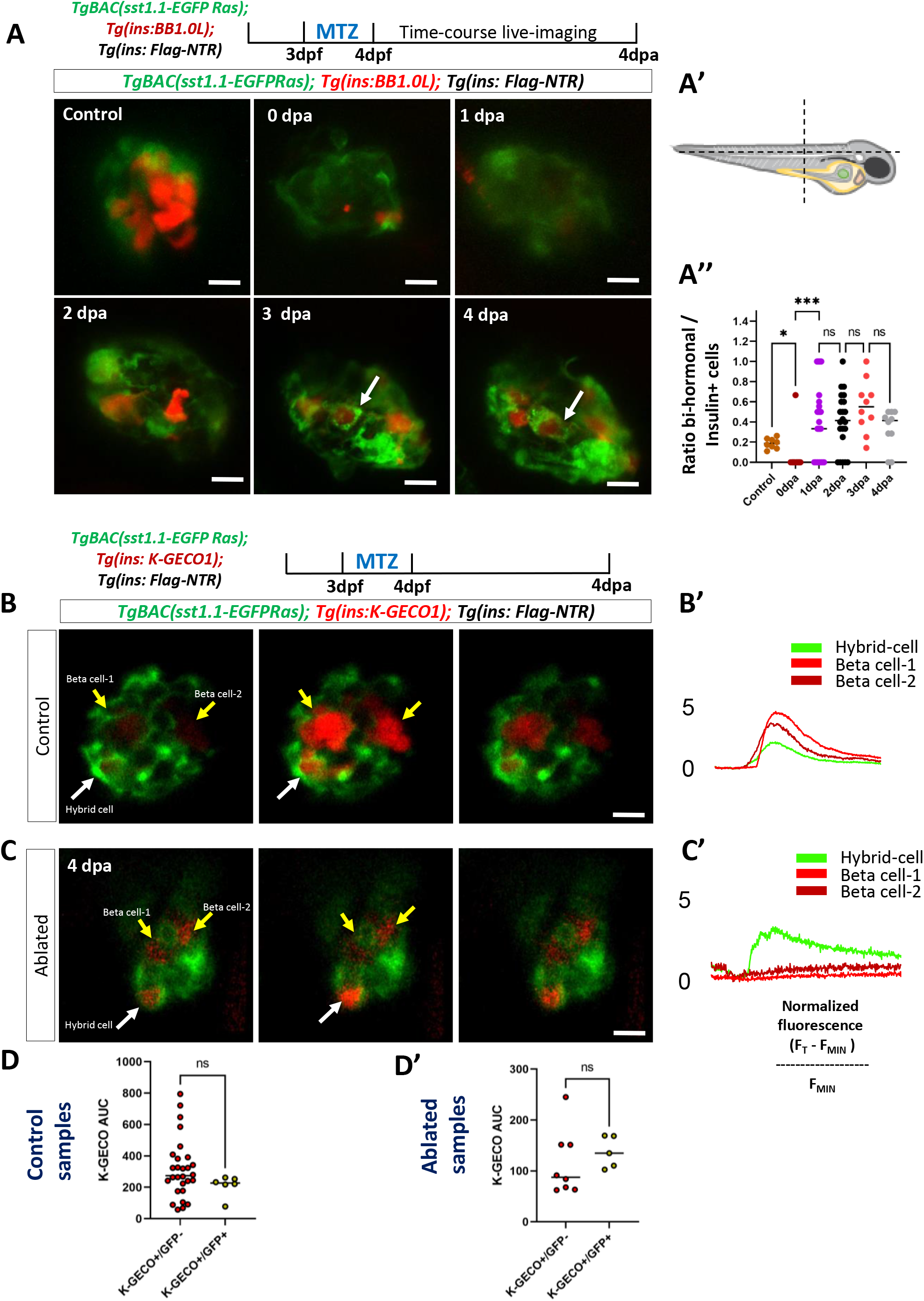
Live imaging of regeneration reveals that the hybrid cells arise *de novo* and acquire glucose-responsiveness. **(A)** To visualize the dynamics of bi-hormonal cells during β-cell regeneration, *Tg(ins: FLAG-NTR)*; *Tg(sst1.1: EGFP-Ras); Tg(ins: BB1.0L)*-triple transgenic animals (5 dpf) were treated with vehicle or Mtz for 24 hours. Following Mtz-mediated ablation of β-cells, a time-course live imaging was performed from 0 to 4 dpa. **(A)** 3D-projections of the zebrafish islet in control and at different time-points post β-cell ablation in the same larva. Following Mtz addition, all tdTomato-expressing cells were ablated and no longer present in the islet. New tdTomato- and GFP-positive cell appear in the islet over time, indicating a *de novo* origin (arrows). **(A’)** Orientation of the zebrafish islet during the live-imaging process. **(A’’)** Quantification of the proportion of bi-hormonal cells during the regeneration process (Brown-Forsythe and Welch ANOVA). **(B, C)** Images from the time-lapse recording of glucose-stimulated calcium influx (6 frames/sec, single plane) of the zebrafish islet in control (Unablated) and at 4-days post β-cell ablation (Ablated). **(B’, C’)** Normalized *K-GECO* fluorescent traces, showing glucose-stimulated calcium influx of bi-hormonal and mono-hormonal cells. **(D, D’)** Quantification of changes in normalized *K-GECO* fluorescence intensity (recorded over 400 frames after glucose-injection) based on area under the curve (AUC) for single-cells. **(D)** Scatterplot for *K-GECO*+/GFP- and *K-GECO*+/GFP+ from unablated samples (Mann-Whitney unpaired t-test, p-value = 0.0907, non-significant). **(D’)** Scatterplot for *K-GECO*+/GFP- and *K-GECO*+/GFP+ from ablated samples (Mann-Whitney unpaired t-test, p-value = 0.222, non-significant). Scale bar: 5µm

Furthermore, we wanted to assess if pre-existing β-cells, spared from ablation, may undergo de-differentiation to form ins/sst bi-hormonal cells. To this end, this we performed β-cell tracing using *Tg(ins: H2B-mEos2); Tg(ins: FLAG-NTR)* double-transgenic zebrafish larvae. These larvae express a green-to-red photoconvertible protein mEos2 (fused to histone H2B) and NTR specifically in the β-cells. At 3 dpf, we photolabeled all pre-existing β-cells in red, followed by Mtz treatment. At 3 dpa, the majority of the hybrid cells displayed green nuclei and somatostatin expression. This strongly suggests that the bi-hormonal cells do not arise from the pre-existing β-cells in the zebrafish larvae (Fig. S14).

### The newly emerging bi-hormonal cells exhibit glucose-stimulated calcium influx *in vivo*

In order to investigate if the emerging bi-hormonal cells, which arise post β-cell ablation, are glucose-responsive and present a calcium influx upon glucose stimulation, we performed *in vivo* calcium imaging using *Tg(sst1.1:EGFP-Ras); Tg(ins:FLAG-NTR);* and *Tg(ins:K-GECO1)* triple-transgenic larvae. We specifically ablated β-cells by exposing the larvae to Mtz from 3 to 4 dpf and performed *in vivo* calcium imaging at 2, 4- and 6-days post β-cell ablation (Movies 1 and 2). Interestingly, we found that the emerging bi-hormonal cells became glucose-responsive as early as 4 days post-β cell ablation (Fig. 7B-C) (Movie 2). The bi-hormonal cells showed a comparable glucose-responsiveness to GFP-negative but *K-GECO1-*positive cells (Fig. 7D), which likely arise via constitutive developmental neogenesis in larvae. The glucose responsiveness of bi-hormonal cells, however, was lower than that of β-cells in unablated animals, likely because the cells were undergoing maturation. Thus, our results indicated that the emerging bi-hormonal cells acquire responsiveness to glucose stimulus.

### Overexpression of *dkk3b* increases hybrid-cell formation

Our initial analysis of gene-expression of the hybrid cells indicated that the cells are enriched for the expression of *dkk3b*, which is also shared with δ1-cells. Moreover, upon β-cell ablation, *dkk3b* was among the genes that showed a trend of increasing expression within the δ1-cell cluster over time (Fig. S8). To test if *dkk3b* plays a role in hybrid-cell formation, we generated transgenic lines with constitutive expression of *dkk3b* under the *insulin* promoter, or with inducible expression based on the *heat-shock promoter* (hsp). In both cases, we observed that the upregulated expression of *dkk3b* in the islets let to increased proportions of insulin and somatostatin double-positive cells during development (Fig. 8A,B). Thus, the forced overexpression of *dkk3b*, which is normally enriched in hybrid and δ1-cells, was sufficient to promote the formation of bi-hormonal cells without injury.

**Figure 8.**
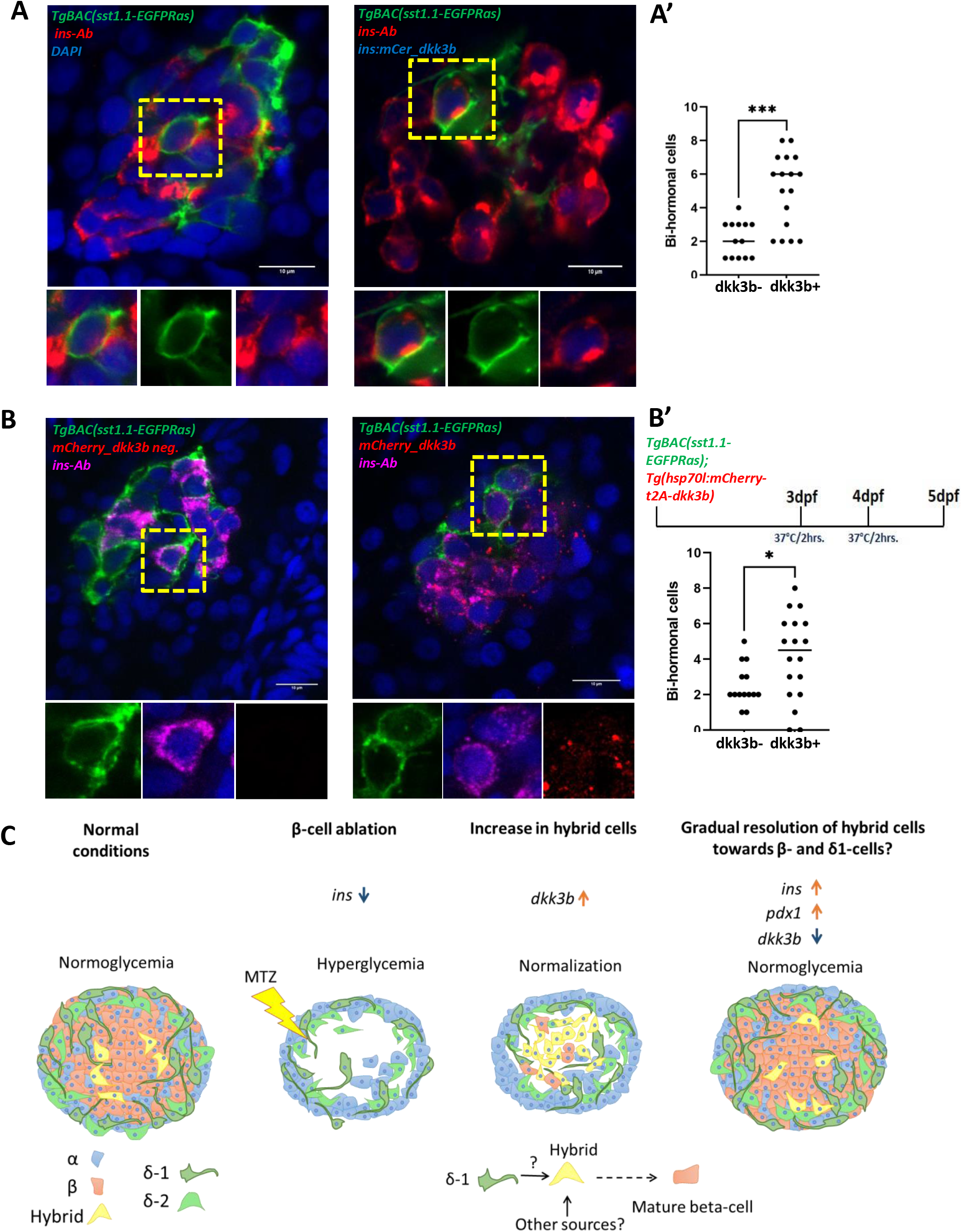
Over-expression of *dkk3b* leads to increased formation of hybrid cells. **(A)** Single-plane confocal images of islets from *Tg(ins: mCerulean_dkk3b)*; *Tg(sst1.1: EGFP-Ras)*-double transgenic animals (5 dpf) stained with insulin antibody (red). The yellow dashed-box outlines the *sst1.1*-*insulin* double-positive cells. **(A’)** Scatter plot showing the number of bi-hormonal cells in wild-type and *ins:mCerulean-t2a-dkk3b* animals (Mann-Whitney unpaired t-test, p-value = 0.0005, significant). **(B)** Single-plane confocal images of islets from *Tg(hsp70l:mCherry-t2a-dkk3b)*; *Tg(sst1.1: EGFP-Ras)*-double transgenic animals stained with an insulin antibody (magenta). The larvae were given a heat-shock for 2 hours at 37°C at 3dpf and 4dpf, thereby inducing the expression of *dkk3b*. The yellow dashed-box outlines the *sst1.1-insulin* double positive cells. **(B’)** Scatter plot showing the number of bi-hormonal cells in wild-type and *Tg(hsp70l:mCherry-t2a-dkk3b)* larvae (Mann-Whitney unpaired t-test, p-value = 0.0352, significant). **(C)** Cartoon model based on the experimental observations in this study. Scale bar: 10µm

### Zebrafish δ1-cells are partially related to γ-cells in mammals

To identify the cellular counterparts of zebrafish δ1- and δ2-cells in the mammalian pancreas, we compared our single-cell transcriptomic dataset (Salem et al., 2019) to human pancreatic data (Segerstolpe et al., 2016). Based on the expression of marker genes in the human dataset, the cells were clustered into α-cells (*GCG*), β-cells (*INS*), δ-cells (*SST*), ε-cells (*GHRL*), γ-cells (*PPY*), ductal (*CFTR*) and acinar (*PRSS1*) cells. The interspecies comparison revealed that zebrafish pancreatic cells clustered with their respective human counterparts, highlighting transcriptomic similarity (Fig. 9A). Notably, we observed that zebrafish ẟ2-cells clustered near to human ẟ-cells, whereas the ẟ1-cells showed partial overlap with human γ-cells. Moreover, there was a higher number of co-expressed genes between zebrafish δ2-cells and human δ-cells, as compared to δ1-cells (Fig. 9B and Table S5). Finally, zebrafish δ1-but not δ2-cells express *pyyb*, encoding peptide YY, a peptide that is also expressed in mouse γ-cells (Perez-Frances et al., 2021) (Table S2 and see Discussion). This analysis suggests that δ1-cells share transcriptional similarity to γ-cells in mammals.

**Figure 9:**
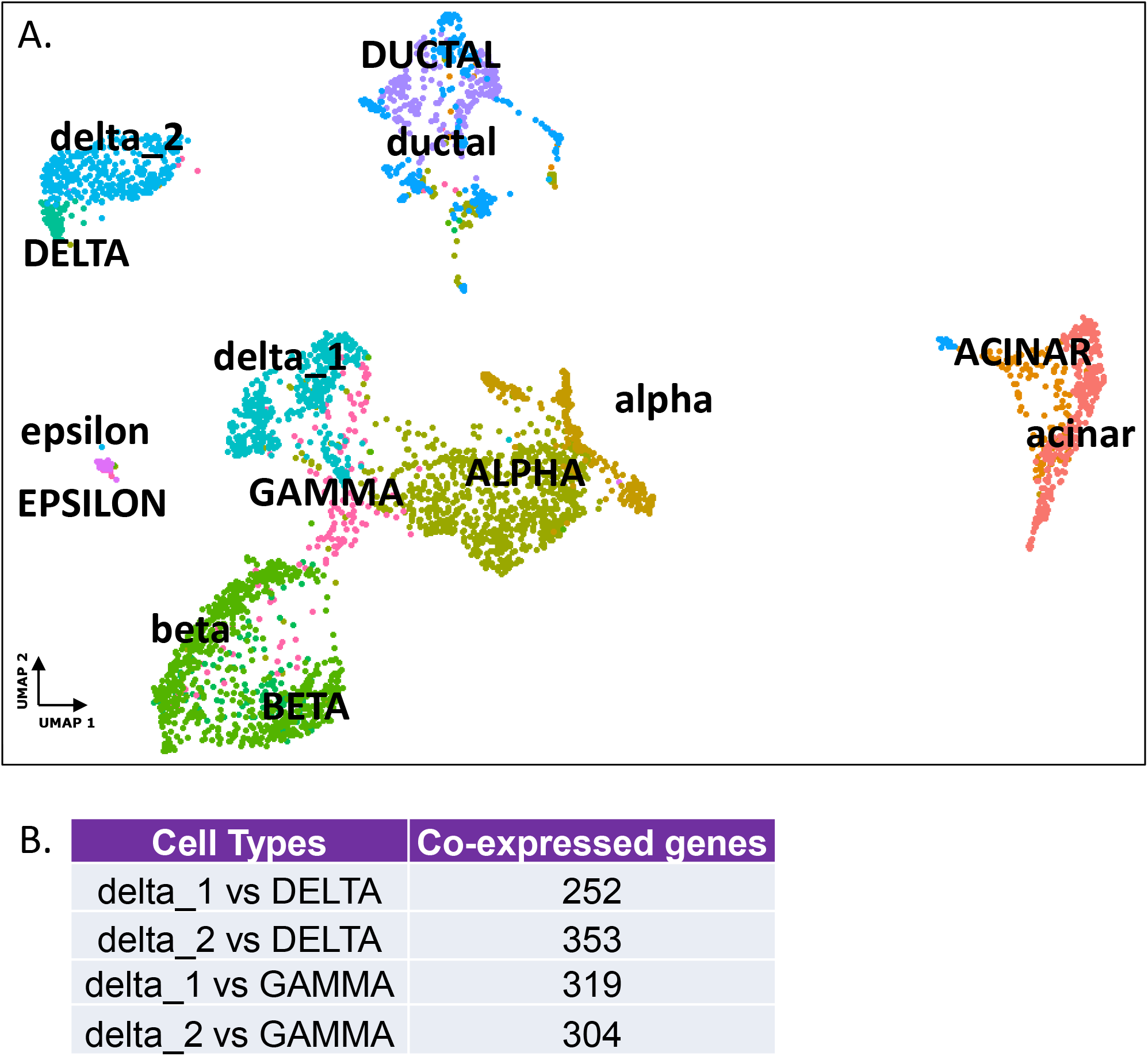
δ1-cells are partially related to human γ-cells. **(A)** UMAP-plot representation of integrated human and zebrafish datasets. Capital case denote human cell types and small case zebrafish cell types. **(B)** Table highlighting the number of co-expressed genes for zebrafish and human cell types.

## Discussion

In this study, we utilized single-cell sequencing to characterize cellular dynamics in a regeneration-competent animal model – the zebrafish (Fig. 8C). Using this approach, we defined the gene-expression of different pancreatic cells upon β-cell destruction and established an atlas of β-cell regeneration, spanning the period from β-cell destruction to emergence of new insulin-expressing cells. Among the different cell types, we characterized a novel population of δ-cells, expressing the *sst* paralogue *sst1.1*, which is consistent with previous studies showing its expression in the zebrafish pancreas (Dalgin et al., 2011; Devos et al., 2002; Spanjaard et al., 2018). In addition, the single-cell resolution of our analysis allowed us to pin-point a small population of bi-hormonal cells, which exist under a hybrid β/δ1-cell identity.

### Hybrid cells as an intermediate hormonal cell population

We show that the hybrid cells share the hormones, fate-determinants and glucose responsiveness of both δ1- and β-cells. While the cells are transcriptionally distinct from β-cells, they show glucose-stimulated Ca^2+^ influx. We validated the presence of hybrid cells at different stages of zebrafish life: from 4 dpf to 16 mpf. Moreover, using single-cell RNA-Sequencing, we compared the transcriptional profile of the hybrid cells to mono-hormonal β-cells and δ1-cells. The differential gene expression analysis revealed an overlapping expression of genes involved in both β and δ-cell fate-specification and function. For example, the cells exhibit intermediate expression of genes involved in insulin prohormone processing, as compared to β-cells. Moreover, these cells express at intermediate levels transcription factors that differentiate between the β- and δ-cell lineages, such as the evolutionary conserved homeobox *mnx1* and *hhex*, respectively (Dalgin et al., 2011; Pan et al., 2015; Zhang et al., 2014). In parallel-unpublished work aimed to define the lineage decisions in the developing endoderm, we find that the population of *sst1.1/ins* cells becomes evident as early as the embryonic islet forms, maintaining similar proportions as in the mature adult islet (∼8% of the insulin-expressing cells) (M.K., unpublished). Moreover, both *sst1.1* and *dkk3b* genes were enriched in the β-cell precursor cluster in a previously published dataset from the embryonic pancreas (Lu et al., 2019). Together, the deep-sequencing experiments suggest that the hybrid cells are an intermediate-lineage population.

### Hybrid cells in rapid resolution of diabetes in zebrafish

Polyhormonal endocrine cells have been observed during embryonic pancreas development in mammals and during regeneration (De Krijger et al., 1992; Teitelman et al., 1993). However, it remains unknown if such cells play a significant role in the pancreas. To test their role, we performed two parallel experiments. First, we asked if the hybrid cells are glucose-responsive. We found that under steady state, the cells behave similar to β-cells, as they exhibit a robust glucose-stimulated calcium influx. Moreover, during regeneration, the emerging hybrid cells acquire the capacity to undergo calcium influx in response to glucose-stimulation. The functional role of the cells in the context of regeneration is supported by the second experiment we carried out: to follow the time course of regeneration using single-cell transcriptomics and to correlate insulin-expression with reversal of diabetes. While there was no evidence for the presence of a major group of regenerated mono-hormonal β-cells even 14 days after β-cell destruction, by this time, the fasting glucose was back to normal, with insulin being expressed predominately in the hybrid cells. Moreover, there was increased expression of *pdx1* and *pcsk1*, which is in line with them acting to restore the islet’s insulin-producing function. Therefore, we show that the hybrid cells fulfill a role in glucose-regulation in the absence of β-cells.

### Hybrid cells inform on islet-cell plasticity

An important unanswered question remains as to the cellular sources of the hybrid cells during regeneration. Our observations point towards the subpopulation of *sst1.1*-expressing cells, which we suspect may increase insulin production upon β-cell destruction, thereby forming hybrid cells. Intriguingly, in mice, δ-cells have been shown to be plastic in juvenile animal but lose their plasticity in adulthood (Chera et al., 2014). It is possible that adult zebrafish maintain an active population of δ1-cells, which covert into hybrid cells and make it possible to restore normoglycemia. However, we cannot exclude the possibility that the centroacinar subpopulation of terminal duct cells, also undergo a process of endocrine differentiation, passing through a bi-hormonal state, which may further resolve into β-cells over time. Differentiating between these two sources will required the development of tools for lineage tracing of δ1-cells in the zebrafish pancreas.

A second important question will be to define the signaling pathways that promote the formation and resolution of such hybrid cells. Here we show that hybrid cells express *dkk3b*, and its overexpression is sufficient to generate hybrid cells in the absence of injury. Since *dkk3b* is predicted to be a secreted factor, the precise cell population that responds to its overexpression to generate hybrid cells remains to be defined. *dkk3b* was previously shown to be expressed in an unknown group of islet cells in zebrafish (Untergasser et al., 2011), and we now reveal its association with the *sst1.1*-expressing δ-cells. Moreover, the human DKK3 is one of the first examples of a heterogeneously expressed gene in islets (Hermann et al., 2007). Notably, *DKK3* is also upregulated in islets from donors with type 2 diabetes (Solimena et al., 2018). The role of this interesting gene in the context of diabetes remains to be studied.

### Hybrid cells and the evolution of endocrine cell types

Our study also raises a curious question regarding the evolution of the islet cells types. Specifically, our interspecies comparison of gene expression illustrated that δ1-cells in zebrafish may be partially-related to γ-cells. In mouse, γ-cells are marked by the expression of *Pyy* together with *Ppy*, which arose from the tandem gene-duplication of the *Pyy gene* (Perez-Frances et al., 2021; Hort et al., 1995). Similarly, we found that zebrafish δ1-cells also expressed *pyyb*. Thus, it is possible that the δ1-cells in zebrafish represent the evolutionary counterpart to γ-cells in mammals. A comprehensive review by Heller noted that the first examples of an islet-like organ containing three distinct endocrine cell types emerged in cartilaginous and bony fish. Later, amphibians and reptiles evolved islets with four major endocrine hormones, and in some but not all mammalian species, islets with all five cell types can be observed (Heller, 2010). We propose that the establishment of hybrid-cell populations may represent an intermediate stage along the diversification and evolution of the discrete subsets of endocrine cells.

Finally, one of the take-home messages is that regenerating imperfect yet functional hybrid cells capable of insulin-production may offer the opportunity to elicit glucose control. Perhaps, such surrogate cells would be stealthy in terms of escaping immune cell recognition and destruction in type 1 diabetes, as they exist under a hidden identity, resembling two different hormonal cell types at the same time.

## MATERIALS AND METHODS

### Zebrafish strains and husbandry

Wild-type or transgenic zebrafish of the outbred AB, WIK, or a hybrid WIK/AB strain were used in all experiments. Zebrafish were raised under standard conditions at 28°C. Animals were chosen at random for all experiments. Published transgenic strains used in this study were *Tg(ins:BB1.0L; cryaa:RFP)* (Singh et al., 2017), *Tg(sst1.1:EGFP-Ras)^fr40Tg^* (Löhr et al., 2018), *Tg(fabp10:dsRed, ela3l:GFP)^gz12^* (Wan et al., 2006), *Tg(ins:FLAG-NTR)*^s950^ (Andersson et al., 2012), *Tg(ins:YFP-2A-NTR3,cryaa:mCherry)^tud201Tg^* (Singh et al., 2017), and *Tg(sst2:RFP)* (Li et al., 2009). Experiments were conducted in accordance with the Animal Welfare Act and with permission of the Landesdirektion Sachsen, Germany (permits: TVV38/2015, TVV 45/2018, TVV33/2019, TVV 32/2020 and all corresponding amendments)

### Single-cell suspension of zebrafish pancreatic islets and liver cells

Single cell suspension of zebrafish pancreas was performed according to the protocol outlined previously (Singh et al., 2018). In brief, primary islets and substantial surround tissue were dissected and dissociated into single cells by incubation in TrypLE (ThermoFisher, 12563029) with 0.1% Pluronic F-68 (ThermoFisher, 24040032) at 37 °C in a benchtop shaker set at 450 rpm for 30 min. During the dissection, we used the fluorescent signal of beta-cells in the *Tg(ins:BB1.0L; cryaa:RFP)* or *Tg(ins:YFP-2A-NTR3,cryaa:mCherry)* reporters in order to orient ourselves in respect to the anatomical position of the main pancreatic islets. Following dissociation, TrypLE was inactivated with 10% FBS, and the cells pelleted by centrifugation at 500g for 10min at 4 °C. The supernatant was carefully discarded and the pellet re-suspended in 500 uL of HBSS (without Ca, Mg) + 0.1% Pluronic F-68. To remove debris, the solution was passed over a 30 µm cell filter (Miltenyi Biotec, 130-041-407). To remove dead cells, calcein violet (ThermoFisher, C34858) was added at a final concentration of 1 µM and the cell suspension incubated at room temperature for 20 minutes. The single cell preparation was sorted with the appropriate gates for identification of β-cells and alive cells (calcein+). FACS was performed through 100 µm nozzle.

For samples from β-cell regeneration time-points, the adult fish were treated with Mtz (Sigma-Aldrich, M3761) which was dissolved at 10mM concentration in fish water. The animals were treated for 24 hours, protected from light. After 24h, Mtz solution was removed and the animals were placed 3 times in fresh fish water before returning them to standard maintenance conditions. Both untreated and Mtz-treated animals were euthanized by an overdose of tricaine. Samples for each time-point comprised of 6 animals (3 males and 3 females). Single-cell suspension was prepared in the same manner (as highlighted above) and 15000 live-cells were FACS sorted for 10X Genomics.

### Single-cell RNA-Seq. of the zebrafish pancreas

The sorted cells (38 µl) were carefully mixed with reverse transcription mix before loading the cells on the 10X Genomics Chromium system (Zheng et al., 2017) in a Chromium Single Cell B Chip and processed further following the guidelines of the 10x Genomics user manual. In short, the droplets were directly subjected to reverse transcription, the emulsion was broken and cDNA was purified using Silane beads. After the amplification of cDNA with 11 cycles, it underwent purification and quantification. The 10X Genomics single cell RNA-seq library preparation - involving fragmentation, dA-Tailing, adapter ligation and a 13 cycles indexing PCR – was performed based on the manufacturer’s protocol. After quantification, the libraries were sequenced on an Illumina Novaseq6000 in paired-end mode (R1: 29 cycles; I1: 8 cycles; R2: 93 cycles), thus generating ∼50-150 mio. fragments. The raw sequencing data was then processed with the ‘count’ command of the Cell Ranger software (v4.0.0) provided by 10X Genomics with the option ‘--expect-cells’ set to 7500 (all other options were used as per default). To build the reference for Cell Ranger, zebrafish genome (GRCz11) as well as gene annotation (Ensembl 95) were downloaded from Ensembl and the annotation was filtered with the ‘mkgtf’ command of Cell Ranger (options: ‘--attribute=gene_biotype:protein_coding -- attribute=gene_biotype:lincRNA –attribute=gene_biotype:antisense’). Genome sequence and filtered annotation were then used as input to the ‘mkref’ command of Cell Ranger to build the appropriate Cellranger Reference.

### Analysis of single-cell RNA-Seq of the zebrafish pancreas

We re-analyzed the single-cell RNA-Seq of the zebrafish pancreatic cells, which was generated using the 10x Chromium pipeline and previously published (Salem et al., 2019). The data is deposited at the GEO repository with an accession number GSE123662. For clustering and gene expression analysis, we utilized Seurat and followed the recommended analysis pipeline. Briefly, the raw data as UMI-counts was log-normalized, regressed to remove the effect of library size, and scaled. Highly variable genes were identified for PCA analysis and graph-based clustering. Marker genes identified for each cluster were used to classify the exocrine, endocrine and ductal population. The endocrine cluster was isolated and sub-clustered to identify and label individual endocrine populations. The cells were filtered using the following QC metrics: 3000 < nFeature_RNA > 350, 25000 < nCount_RNA > 2500 and percent.mito < 20. All the cells that passed the quality check were used for downstream analysis.

To define the changes the abundance of δ1-cells during regeneration, at each time point, we calculated the ratio between the number of *δ1-cells* and α-cells, filtering out any clusters that expressed multiple hormones apart from the combination of *sst1.1* and *ins*. We calculated 99% confidence intervals from a 1000-fold random sampling of the total number of cells present at every time from a binomial distribution with the observed cell type frequencies as probabilities, using the function rbinom in base R.

### Comparison of zebrafish and human datasets

To identify transcriptional overlap between cell-types from human and zebrafish pancreas, we quantified the number of co-expressed genes between cell clusters from single-cell dataset of zebrafish (Salem et al., 2019) and human (Segerstolpe et al., 2016). We converted the human genes to their corresponding zebrafish homologues using the mapping from bioMart. For human genes with multiple zebrafish orthologues, the gene was mapped to each orthologue. Next, we integrated the modified human dataset and the zebrafish dataset using ScTransform function in Seurat. We used the “scaled.data” function to identify co-expressed genes. The mean of the scaled expression was calculated for all genes in a specific cluster. The genes with the mean greater than 0 were considered as being expressed.

### ATAC-Seq. of zebrafish β-cells and hepatocytes

ATAC-Seq. was carried out using 50,000 cells collected using FACS. For FACS, single-cell suspension was generated as mentioned above. For ATAC-Seq. of β-cells, tissue for dissociation was collected from *Tg(ins:BB1.0L; cryaa:RFP)* line and β-cells sorted using the red fluorescence. Cells were processed according to the protocol outlined in (Buenrostro et al., 2013). In brief, cells were homogenized in a 2 ml Dounce homogenizer with 5 strokes in 1X homogenization buffer on ice. The nuclei suspension was centrifuged at 500g for 10 min at 4 °C. After removing the supernatant, transposition was carried out by incubating the nuclei in the transposition reaction mix for 30 min at 37 °C. Transposed DNA was purified using Qiagen MinElute Reaction Cleanup Kit. Barcoded libraries were generated from the isolated DNA by PCR amplification for 11 cycles. Libraries were sequenced on llumina HiSeq2500 in 2 × 75 bp paired-end mode. We conducted two biological replicates for β-cells. Processing of ATAC-seq. raw data was done according to the pipeline from the Kundaje lab (Koh et al., 2016). MACS2 (Feng et al., 2012) was used to identify the peak regions. Only peaks present in both the biological replicates of β-cells were retained. The β-cell specific peaks were exported as a bed file and visualized in IGV viewer (Robinson et al., 2011).

### Transcriptional profiling and analysis of β-, δ1-, and hybrid cells

Collection of β-, δ1-, and hybrid cells for transcriptional profiling was performed using FACS of single-cell suspension of dissected islets from *Tg(ins:BB1.0L); Tg(sst1.1:EGFP-Ras)*. For single-cell profiling using Smart-Seq., cells were collected in 96-well plate using index sorting and processed according to the previously published pipeline (Singh et al., 2018). Analysis was performed using Seurat (Satija et al., 2015), with cells classified using their position on the FACS plot (δ1-cells: Green+, Red-; hybrid cells: Green+, Red+; and β-cells: Green-, Red+). For bulk-sequencing and analysis, 2,000 cells of each type were collected in a 1.5 ml tube and processed according to the pipeline published in (Janjuha et al., 2018). Differential gene expression analysis was analyzed using DESeq2 (Love et al., 2014).

### Pseudotemporal analysis of single-cell RNA-seq. data

Pseudotemporal analysis was performed using Monocle 2 (Trapnell et al., 2014) following the recommended analysis protocol (http://cole-trapnell-lab.github.io/monocle-release/docs/). Briefly, raw count data was imported into a Monocle object, normalized using the estimated size factors, and filtered to remove low quality genes and cells. Gene dispersion was estimated using negative binomial distribution with fixed variance, and used to identify the highly variable genes in an unbiased manner for pseudotemporal ordering. Pseudotemporal trajectory was constructed using the ‘DDRTree’ algorithm, and plotted on a 1-D graph with jitter to enhance visualization. Cells in the graph were colored according to their FACS plot identity. Gene Expression dynamics in relation to pseudotime was calculated using the command ‘differentialGeneTest (Monocle_object, fullModelFormulaStr = “∼sm.ns(Pseudotime)”)’, and plotted using the heatmap function for visualization.

### Diffusion pseudotime analysis of single-cell RNA-seq. data

For single cell trajectory inference, diffusion pseudotime was computed for 7 dpa and 14 dpa and solely on selected cells (*ins*+ and *sst1.1*+ cells) following the standard analysis pipeline (Haghverdi et al., 2016) provided by Scanpy package. Since a single lineage of a timepoint was selected, the number of branchings was set to 1. For cell trajectory analysis, diffusion maps were developed. For 14 dpa, the *sst1.1*-high cells were selected as root cells to capture the transition of δ1-cells. To visualize the marker genes trajectory on trendline plot, plot.wishbone_marker_trajectory function was used provided by the Wishbone package (Setty et al., 2016).

### Tissue collection for imaging

To facilitate confocal imaging of the islets, the pancreas was dissected from fish (larval) or the gut (juvenile and adults) after fixation. Fish were killed in Tricaine prior to either direct fixation or dissection of gut, and the samples immersed in 4% paraformaldehyde + 1% Triton-X for 2 days at 4 °C. The pancreas was then manually dissected and washed multiple times in PBS.

### Immunofluorescence and image acquisition

Whole-mount immunofluorescence was performed on pancreas collected as described above. The collected samples were permeabilized in 1% PBT (Triton-X-100) and blocked in 4% PBTB (BSA). Primary and secondary antibody staining were performed overnight at 4 °C. Primary antibodies used in this study were anti-insulin (guinea pig, Dako A0564)/insulin (rabbit, Genetex 128490) at 1:200, anti-somatostatin (rat, Genetex GTX3906) at 1:200, and anti-EGFP (chicken, Abcam ab13970) at 1:250. Secondary antibodies at 1:500 dilutions used in this study were Alexa Fluor 568 anti-guinea pig, Alexa Fluor 568 anti-rabbit, Alexa Fluor 647 anti-rabbit and Alexa Fluor 647 anti-rat. Samples were mounted in Vectashield and imaged using a Zeiss LSM 780. ImageJ was used to add scale bars and PowerPoint was used for adding arrows and labels.

For immuno-staining of the adult pancreatic sections obtained from controls and various time-points post β-cell ablation, the Tyramide Signal Amplification kit (Thermofisher, Alexa Fluor™ 555 Tyramide SuperBoost™ B40923) was used to amplify the signal from the insulin antibody. Primary and secondary antibody staining were performed as recommended by the kit. Draq7 was used as the nuclear label. Sections were mounted in Vectashield and imaged using Zeiss LSM 780.

### Live-imaging during β-cell regeneration

From 1 dpf onwards, *Tg(ins:FLAG-NTR)*; *TgBAC (sst1.1:EGFP Ras), Tg (ins: BB1.0L)*-triple transgenic zebrafish larvae were treated with 0.003% 1-phenyl thio-urea (PTU) (200µM) to inhibit pigmentation. For β-cell ablation, 3 dpf larvae were incubated with 10 mM Mtz (Sigma-Aldrich, M3761) dissolved in E3 medium containing 1% DMSO, and maintained for 24 h in the dark before they were rinsed and returned to fresh E3 medium. E3 medium with 1% DMSO was used as control. Following the ablation, individual larvae was mounted in a glass-bottom micro-well dish (MakTek Corporation) in 1% low-melting agarose (LMA) containing 0.4g/litre tricaine. After the agarose was solidified, the dish were filled with E3 medium containing 0.4g/litre tricaine. Live-imaging was performed on an upright laser-scanning confocal microscope, Zeiss LSM 780. EGFP and dsRed signals were acquired simultaneously using the 488nm and 561 nm laser lines. Laser power was maintained as low as possible to minimize photo-toxicity.

### Quantification of bi-hormonal cell positions in the islet

Pancreatic sections from 7 and 16 mpf *Tg(sst1.1: EGFP-Ras)* transgenic zebrafish were co-stained with insulin and somatostatin antibodies. The number of bi-hormonal cells was quantified manually using ImageJ software. The islet periphery was determined using the “freehand” selection tool and the islet center was computed using the “center of mass” measurements in ImageJ (Analyze -> Set Measurements -> Center of Mass). The relative position of each bi-hormonal cell was determined in the following way: relative distance = (distance from the islet center) / (distance from the islet center to the islet edge nearest to the bi-hormonal cell). The relative position of bi-hormonal cells was represented as a scatter-plot with the position of the cells on the x-axis (islet center = 0.0; islet periphery = 1.0) and the age of the fish on the y-axis.

### β-cell tracing in zebrafish larvae

From 1 dpf onwards, *Tg(ins: H2B-mEos2; cryaa: CFP)*; *Tg(ins: FLAG-NTR; cryaa: mCherry)* double transgenic zebrafish larvae were treated with 0.003% PTU (200µM) to inhibit pigmentation. At 3 dpf, the zebrafish larvae were anaesthetized using tricaine and exposed to blue light for photoconversion. From 3 to 4 dpf, the larvae were incubated in 10 mM Mtz dissolved in E3 medium containing 1% DMSO and maintained for 24 hours in the dark. Mtz was washed and the fish were returned to fresh E3 medium.

### *In vivo* calcium-imaging in zebrafish larvae

Live-imaging and glucose-injections were performed as described previously (Salem et al., 2019)

### Generation of Tg(ins:nls-mCerulean-T2A-dkk3b; cryaa:mCherry) zebrafish line

The dkk3b coding sequence was purchased from Zebrafish Gene Collection (ZGC) cDNA clones (MGC: 162332). The cDNA was PCR amplified to flank the cDNA with SpeI/PacI restriction sites. The PCR product was digested with SpeI/PacI restriction enzymes. The ins:nls-mCerulean*; cryaa:mCherry* construct (used for generating *Tg(ins:nls-mCerulean; cryaa:mCherry)* line) was digested with SpeI/PacI. The two DNA fragments were ligated and transformed to generate ins:nls-mCerulean-T2A-dkk3b; cryaa:mCherry plasmid. The entire construct is flanked with I-SceI sites to facilitate transgenesis. I-SceI based transgenesis was carried out, and founders selected based on mCherry (red) fluorescence in the eye, and mCerulean (blue) expression in the pancreatic islet.

### Generation of Tg(Tol2-hsp70l:mCherry-T2A-dkk3b) zebrafish line

The dkk3b coding sequence was PCR amplified to flank the cDNA with NheI/BglII restriction sites. The PCR product was digested with NheI/BglII restriction enzymes and was fused to mCherry using the T2A sequence. Thereafter, mCherry-t2a-dkk3b sequence was cloned downstream of the hsp70l promoter region. The entire construct is flanked with Tol2 sites to facilitate transgenesis. Tol2 based transgenesis was carried out, and founders selected based on ubiquitous expression of mCherry (red) fluorescence in the zebrafish larvae upon heat-shock at 37 degrees.

### Generation of Tg(ins:K-GECO1; cryaa:mCherry) zebrafish line

For the construction of the *Tg(ins:K-GECO1; cryaa:mCherry)* line, we used PCR amplification with primers designed to introduce 5’ EcoRI and 3’ PacI restriction enzymes sites in the cDNA for K-GECO1. The established plasmid backbone containing ins:mAG-zGeminin; cryaa:mCherry (Ninov et al., 2013) was digested with EcoRI/PacI and K-GECO1, and the cDNA was ligated using the EcoRI/PacI sites. The construct was flanked with I-SceI sites to facilitate transgenesis. Several founders were screening and a founder with Mendelian segregation were selected for expanding the transgenic line.

### Generation of Tg(ins: H2B-mEos2b; cryaa: CFP) zebrafish line

For the construction of the *Tg(ins: H2B-mEos2b; cryaa: CFP)* transgenic line, the cDNA for H2B-mEos2b was PCR amplified to flank the cDNA with Mfe1/Pac1 restriction enzyme sites. A previously established plasmid backbone with ins: mKO2-zCdt1; cryaa: CFP (Ninov et al., 2013) was digested with EcoRI/PacI and H2B-mEos2b was ligated using the EcoRI/PacI sites (Mfe1 and EcoR1 have compatible cohesive ends). The entire construct is flanked with I-SceI sites to facilitate transgenesis. I-SceI based transgenesis was performed and founders were selected based on CFP (blue) fluorescence in the eye, and green expression in the pancreatic islet.

### Electron Microscopy

Correlative Light and Electron Microscopy (CLEM) of immunolabeled resin sections was performed as previously described (Fabig et al., 2012). In brief, WT and ablated zebrafish larvae were embedded in Lowicryl K4M using the progressive lowering of temperature method (Carlemalm et al., 1982). Ultrathin resin sections were cut on Leica UC6 ultramicrotrome (Leica Microsystems Wetzlar, Germany), labeled with primary anti-GFP (rabbit-anti-GFP, TP401 from Torrey Pines) followed by consecutive incubations with protein A 10 nm gold, goat-anti-rabbit Alexa 488, and DAPI. Sections were analysed on a Keyence BZ 8000 fluorescence microscope, contrasted with uranyl acetate, dried, and imaged on a Jeol JEM 1400-Plus transmission electron microscope (Jeol, Freising, Germany) running at 80 kV.

### Statistical analysis

Statistical analysis was performed using R and GraphPad Prism. No animals were excluded from analysis. Blinding was not performed during analysis. Analysis of normal distribution was not performed. To compare the proportion of *sst1.1* expressing β-cells during regeneration, an unpaired two-tailed t-test with unequal variance (t-test(x = dataframe, alternative = “two.sided”, paired = FALSE, var.equal = FALSE)) was used. A p-value of <0.05 was considered statistically significant. For plotting, Tukey style boxplots showing the median along with the 25th percentile (lower quartile -Q1) to 75th percentile (upper quartile - Q3) range were used. For all boxplots, whiskers extend to 1.5 times the interquartile range (IQR = Q3 − Q1). In addition, individual data points are plotted on the graphs. To compare the proportions of bi-hormonal cells during time-course imaging, to evaluate the AUC during in-vivo calcium imaging and the bi-hormonal cell number upon dkk3b over-expression, Mann-Whitney unpaired t-test (non-parametric test) was used for statistical analysis. The graphs were prepared using Graph Pad prism software and p-value of <0.05 was considered statistically significant.

### Data availability

The processed data are available for viewing and exploration on the publicly accessible Single Cell Portal at https://singlecell.broadinstitute.org/single_cell/ study/SCP1549. Single-cell RNA sequencing data of zebrafish pancreatic cells have been deposited in GEO under accession number GSE123662 (Salem et al., 2019). Single-cell and bulk analysis datasets of pancreatic somatostatin and β-cells from zebrafish have been deposited in GEO under accession number GSE152697. Data for ATAC sequencing of zebrafish pancreatic β-cells and hepatocytes have been deposited in GEO under accession number GSE152199.

## Contributions

Conceptualization: S.P.S, N.N; methodology: S.P.S, P.C., A.H, M.K., L.D.S, S.J., O.K., S.B., J.B., A.K., A.P., T.K., S.R.; formal analysis: S.P.S., P.C. L.D.S, B.S., P.O.; S.E.E; resources: N.N, J.P.J.; writing (original draft): S.P.S, N.N.; supervision: J.P.J. and N.N.; funding acquisition: J.P.J. and N.N.

## Acknowledgements

Work in J.P.J.’s laboratory was funded by a European Research Council Starting Grant (ERC-StG 715361 SPACEVAR). We are grateful to M. Hammersmith for sharing *Tg(sst1:GFP-Ras)*. Work by S.P.S. is supported by the Fonds de la Recherche Scientifique-FNRS under Grant(s) n° 34772792 – SCHISM. N.N. receives funding from the Center for Regenerative Therapies Dresden at TU Dresden, the German Center for Diabetes Research (DZD), as well as research grants from the German Research Foundation (DFG), and the International Research Training Group (IRTG 2251): ‘Immunological and Cellular Strategies in Metabolic Disease’. We are grateful to the following facilities at the CMCB and the CRTD: Deep Sequencing, Flow Cytometry Facility, Electron Microscopy, Light Microscopy, Zebrafish.

